# Direct observation of a crescent-shape chromosome in *Bacillus subtilis*

**DOI:** 10.1101/2023.02.09.527813

**Authors:** Miloš Tišma, Florian Patrick Bock, Jacob Kerssemakers, Aleksandre Japaridze, Stephan Gruber, Cees Dekker

## Abstract

Bacterial chromosomes are folded into tightly regulated three-dimensional structures to ensure proper transcription, replication, and segregation of the genomic information. Direct visualization of the chromosomal shape within bacterial cells is hampered by cell-wall confinement and the optical diffraction limit. Here, we combine cell-shape manipulation strategies, high-resolution fluorescence microscopy techniques, and genetic engineering to visualize the intrinsic shape of the bacterial chromosome in real-time in live *Bacillus subtilis* cells. We show that the chromosome exhibits a crescent shape with a non-uniform DNA density that is increased near the origin of replication (*oriC*). Additionally, we localized ParB and BsSMC proteins – the key drivers of chromosomal organization – along the contour of the crescent chromosome, showing the highest density near *oriC*. Opening of the BsSMC ring complex disrupted the crescent chromosome shape and instead yielded a torus shape. These findings help to understand the threedimensional organization of the chromosome and the main protein complexes that underlie its structure.

## Introduction

Over the past decade, it has become evident that bacterial chromosomes are folded into a compact 3D architecture that regulates transcription and is necessary for cell survival ^1–3^. As nuclear compartmentalization is absent in bacteria, a high abundance of DNA-binding proteins can locally bind and change the chromosomal structure ^4^. Furthermore, the chromosome is subject to the crowding effects of cytosolic components ^5^ and the genome has to fit within the confines of the cell boundary which squeezes the millimeter-long bacterial genome within the micron-size cell ^6^. Although bacterial chromosomes have been imaged by microscopy in many earlier works ^7–12^, the tight confinement of the genome combined with the finite optical diffraction limit has been a hindrance towards resolving its intrinsic shape and structure. Attempts to significantly increase the spatial resolution via super-resolution microscopy techniques often require synthetic dyes that include lengthy washing procedures or invasive crosslinking steps which may affect the DNA structure and dynamics in live cells ^7,9,13,14^. Direct high-resolution imaging of the chromosome would shed light onto the chromosomal dynamics, macrodomain organization ^15,16^, and the roles of various proteins in its organization.

An indirect but powerful method of studying the chromosomal structure is provided by chromosome capture techniques (3C^17^, 4C^18^, 5C^19^), most notably ‘Hi-C’^9,20–25^. This technique uses proximity-based ligation in combination with next-generation sequencing to uncover the average spatial organization of the DNA within a population of cells^22^. Over the past decade, Hi-C has been established as a major tool for studying the 3D chromosome organization and it has contributed key insights about the chromosome organization in many bacterial species^9,20,21,24–26^. Interestingly, the most well-studied bacterial model organisms *Escherichia coli* and *Bacillus subtilis* show an entirely different chromosomal structure in Hi-C maps. *E. coli* was deduced to have a textbook circular chromosomal shape with separately condensed chromosomal arms that are twisted within the cylindrical cell ^20^. *Bacillus subtilis* however, shows a distinct “second diagonal” feature in the Hi-C maps ^9^ which indicates that two chromosomal arms are spatially aligned, which presumably is facilitated by the action of bacterial structural maintenance of chromosome (SMC) proteins ^9,24,25^. However, questions remain as Hi-C methods involve extensive cell fixation which can alter the DNA organization ^22^, and conclusions are drawn from population averages where the single-cell structure and cell-to-cell variabilities are lost.

A promising way to spatially resolve the bacterial chromosome is to use cell-shape manipulation techniques ^16,27–30^ or expansion microscopy (ExM) ^31–33^. While expansion microscopy, where bacteria are encapsulated in a hydrogel that is stretched, requires cell fixation that precludes live cell imaging ^33^, cell-shape manipulation techniques allow for imaging of live cells, while a single chromosome can be maintained via DNA replication halting. Using this approach, we previously visualized the *E. coli* chromosome at the single-cell level, showing a toroidal shape with fast local dynamics and interesting substructures within the genome ^16,27^.

Here, we use cell-shape manipulation with replication halting to capture the structure of the *Bacillus subtilis* chromosome in single live cells. Cell-shape manipulation relaxed the cell-boundary confinement of the chromosome in live bacteria, thus allowing standard super-resolution microscopy techniques to capture the intrinsic chromosome shape. We observed that the *B. subtilis* chromosome adopts a crescent shape, with an origin region at one tip of the crescent that is highly condensed. Furthermore, we measured the positioning of the key chromosome-organizing proteins ParB and BsSMC along the contour of the chromosome. Upon BsSMC disruption, the crescent chromosome shape changed into a torus shape, indicating that BsSMC is required for maintaining the crescent shape. The data provide insight into the chromosomal organization of the *B. subtilis* chromosome and its organizing proteins.

## Results

### Cell-shape manipulation of *Bacillus subtilis* bacteria

In standard growth conditions, both for nutrient-rich or in a minimal medium, *B. subtilis* cells grow into rod shapes with its chromosome residing close to the center of the cell as a confined object of ~1.3 μm x 0.7 μm (Fig. S1A-B) ^6^. Due to the diffraction limits of microscopy, most of the chromosome’s inner structure cannot be resolved. To eliminate the confining effects set by the cell wall, we converted the rod-shaped *B. subtilis* cells into spheroidal cells (also known as L-form cells ^34,35^) similarly to the protocol described in Kawai *et al*.^36^ (Figure 1C, *see Methods*). The physical conversion from a capped cylinder to a spheroid implies a change in surface-to-volume ratio. In our case, the total surface remains constant as it is set by the cell membrane that cannot substantially grow over the course of the fast conversion to spheroidal cells. To accommodate the change, we used different osmotic media to promote water uptake into the spheroidal cells, and accordingly a volume increase. We tested a variety of media ranging from highly hypoosmotic (100mM osmolyte) to isoosmotic concentrations (500mM osmolyte, see *Methods*), which did not affect growth in cylindrical cells (Fig. 1C, S1C). Under wall-less conditions, we observed the highest increase in volume under slightly hypoosmotic conditions, viz., 300mM osmolyte (Fig. S1D). Cylindrical cells of, on average, 3.3 × 0.8 × 0.8 μm^3^ size then adopted an approximately spheroidal shape with an average size of 2.3 × 2.3 × 1.7 μm^3^. This resulted in the average total volume change of up to about a factor of 3 of the original volume (Fig. 1D). This volume expansion increased the physical space where the chromosome can freely reside, allowing for observation of its intrinsic shape without confinement.

**Figure 1.**
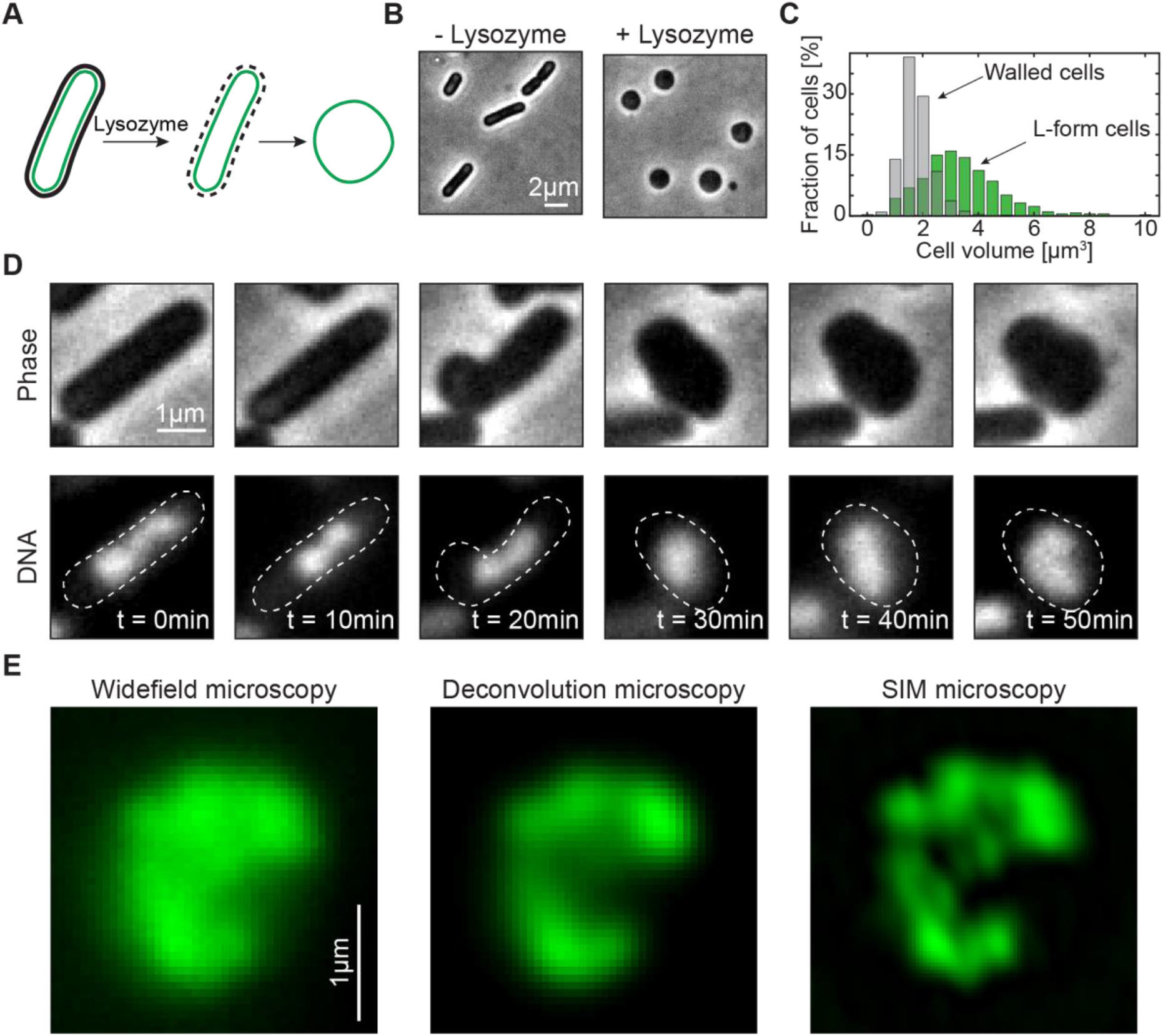
*Bacillus subtilis* chromosome adopts a crescent shape upon cell widening. **A)** Graphical representation of the cell-shape conversion from rod-shaped to spheroidal shape. **B)** Conversion of rod-shaped cells to spheroidal cells under hypoosmotic conditions in SMM+MSM medium (300 mM succinate) upon addition of lysozyme. **C)** Total cell volume before (grey, V_avg_=1.8 ± 0.6 μm^3^, *N=550*) and after the lysozyme treatment (green, V_avg_=3.4 ± 1.5 μm^3^ *N=1056*). **D)** Example timelapse imaging of the conversion of a single cell from rod shape to spherical shape under an agar pad for the BSG217 strain at 37°C. **E)** Comparison of widefield (left), deconvolved (middle), and Structured Illumination Microscopy (SIM) (right) image of a crescent chromosome in the strain BSG217.

### Observation of a crescent chromosome *in Bacillus subtilis* in single cells

To obtain the image of a single chromosome in individual cells, we constructed a strain containing temperature-sensitive DnaB protein (*dnaB134ts*, Fig. S2, Table S1)^10,37,38^. This protein is an essential component during the DNA replication process in *B. subtilis* as it facilitates new rounds of initiation by facilitating helicase loading^39,40^. Upon temperature increase to 45°C, the strain carrying DnaB^K85E^ (*dnaB134ts* locus) experiences a strong inhibition of initiation of new DNA replication rounds^38^. To control for the possible denaturation of HbsU protein at high temperatures ^41^ (which would alter chromosomal structure), we grew the strain at a maximum of 39°C. Additionally, we constructed a strain carrying *P_lac_-sirA* whose protein product can also efficiently halt the initiation of DNA replication by directly inhibiting the action of DnaA^42,43^ (Fig. S2, Table S1), whilst preserving HbsU stability at 30°C ^41^. For DNA imaging, we used a fluorescently labeled version of the HbsU protein, which is a protein that is uniformly bound across the DNA^44,45^, or synthetic DNA intercalating dyes (Fig. S2) that do not affect cell viability ^46^, as well as a standardized DNA dye (DAPI). All different fluorophores and replication-halting variants yielded similar results (Fig. S2).

In a large fraction (~40%) of the cells, we could resolve the intrinsic chromosome shape and observed that the *B. subtilis* chromosome exhibited a crescent shape (Fig. 1D, S3A, B). This was observed irrespective of the chromosome labeling or replication-halting strategy (Fig. S2). In contrast to the observations in *E. coli*, the final chromosome size was smaller than the cell diameter (Fig. S4), which suggests that the chromosomes did largely expand to their intrinsic shape. A considerable cell-to-cell variability was observed where some chromosomes appeared as a more condensed object that we were unable to resolve (Fig. S3B).

Next, we used superresolution microscopy to obtain high-resolution images of single *B. subtilis* chromosomes. Deconvolution of widefield microscopy and structured illumination microscopy (SIM) allowed for the imaging of live cells with spatial resolution down to 150 nm and 120 nm, respectively. Images taken with both techniques consistently showed that chromosomes adopted a crescent shape (Fig. 1E, S5, Movie S1, S2). This is consistent with previous Hi-C data that suggested a close spatial alignment of left- and right-chromosomal arms ^25^. Interestingly, the fluorescence intensity along the contour of the chromosome appeared to be quite variable between two ends as well as from cell to cell. This suggests a dynamic compaction level of the DNA along the chromosome where DNA is distributed non-uniformly (Fig. 1D-E). To quantitatively measure the physical characteristics of the crescent-shaped chromosomes, we ensured the presence of a single chromosome within each cell, as any unfinished replication round could result in large changes in the DNA amount and chromosome shape, thus obstructing quantitative assessments of the chromosome shape, size, and dynamics.

### DNA is highly compacted in the origin region

First, we located the origin of replication (*oriC*) as the reference point. Conveniently, *B. subtilis* has multiple *parS* sites around the *oriC*, which bind partitioning protein B (ParB) in high numbers^47^ (Fig. 2A). ParB proteins bridge multiple *parS* sites and form a single focus per chromosome^48,49^, and they are commonly used as a proxy for the number of chromosomes per cell^25,50–53^, given that newly replicated origins are quickly separated. We used a fluorescent fusion of ParB-mScarlet in addition to *P_lac_-sirA* in order to obtain single chromosomes with a single fluorescently labeled origin-proximal region^54^ (Fig. 2B-E, Table S1). In growth assays, this strain behaved equivalent to wild-type *B. subtilis* (Fig. S6A), and it showed the same phenotype under the microscope (Fig. S6B). Upon induction of SirA protein, up to 78% of the cells showed a single focus per cell, indicating successful replication halting (Fig. 2E). In these cells, we observed the majority of ParB-mScarlet fluorescence to localize at the tip of the crescent chromosome (Fig. 2F, S7).

**Figure 2.**
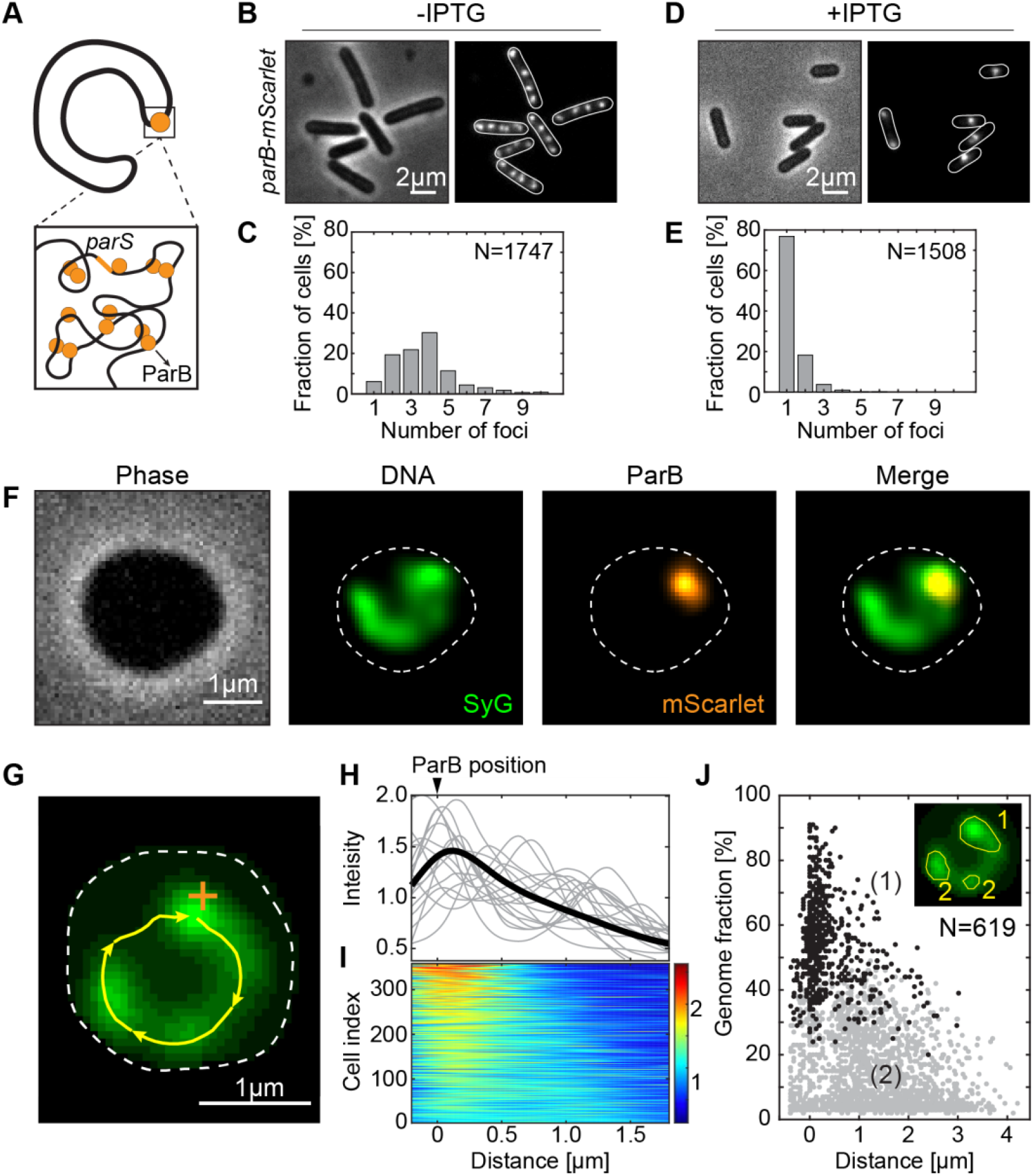
ParB complex co-localizes with a region of high DNA density. **A)** Graphical representation of the crescent-shape chromosome. The zoom sketches the origin/*parS* region that is condensed by ParB proteins (orange). **B)** Phase (left) and fluorescence (right) images of the BSG4595 cells (*parB-mScarlet; P_hyperspank_-sirA*) in the absence of ITPG. **C)** Quantification of the number of fluorescent ParB-mScarlet foci in single cells imaged as in panel B (*N=1747*). **D)** Same as panel B but for cells in the presence of 2 mM ITPG for 150 min. **E)** Quantification of the number of fluorescent ParB-mScarlet foci in single cells imaged as in panel D (*N=1508*). **F)** Representative example of a cell after treatment with lysozyme (400 μg/ml) for 30 min. **G)** Crescent-shaped chromosome. Cell outline is denoted by the white line; DNA contour track by yellow line; location of the ParB focus by the orange cross. **H)** DNA intensity along the contour of the chromosome (cf. yellow line in panel G). Black line shows the average normalized intensity obtained from all cells (N=619). Gray lines display arbitrarily chosen individual examples. The position of the ParB focus is indicated on top (defining the 0 μm position). **I)** DNA density along the chromosome in all individual cells. Colorbar represents the fold-increase. Cells are ordered from top to bottom in terms of contrast. **J)** Cluster analysis of the chromosome. Inset shows an example output of the analysis with 1 primary and 2 secondary-size clusters. The fraction of the genome that is contained within the primary focus (1) (black dots) is plotted versus distance from the origin (ParB focus; n=619). Secondary condensation (2) foci are represented as gray dots.

After thus successfully localizing the *oriC* and ensuring the presence of a single chromosome, we proceeded with the analysis of DNA compaction along the contour of the crescent chromosome (Fig. 2G, see Methods and Ref. 16). In all cells, we observed an increase in the DNA fluorescence intensity proximal to the ParB focus, i.e., at the origin of replication (Fig. 2H-I). Fluorescent intercalating dyes as SyG, bind uniformly to the DNA^55^, thus allowing to correlate the fluorescence intensity with the underlying amount of DNA. We observed an uneven distribution of DNA along the contour of the chromosome, with a higher abundance of DNA present near the origin of replication and less DNA present towards the tail of the crescent chromosome. Using previously established analyses^16^, we detected DNA clusters, representing more condensed regions, along the contour of the chromosome (Fig. 2J). We termed the cluster with the highest amount of DNA as the ‘primary cluster’, and all other clusters found as ‘secondary clusters‘ (Fig. 2J). We found that the primary clusters typically localized near the *ori* region (Fig. 2J – black data), i.e., coinciding with the ParB focus. Most of cells possessed a secondary condensed region, albeit variable in size and position along the chromosome (Fig. 2I – grey data). We did not observe a particularly preferred chromosome position for these secondary clusters, and such domains instead appeared to position at arbitrary locations along the chromosome contour shape as well as contained a variable amount of DNA.

As the primary clusters tended to be positioned near *ori*, we analyzed the total DNA percentage that was localized near *ori* (Fig. S7). We observed that the majority of cells (67%) exhibited a very large fraction (>40%) of their entire genome within only 500nm from the origin of replication. Our data thus show a high DNA condensation proximal to the origin region, and concomitantly lower DNA condensation along the rest of the chromosome.

### BsSMC proteins spread along the entire contour of the crescent chromosome

SMC proteins constitute key players in the large-scale chromosome organization in all domains of life ^56^. For *B. subtilis*, this concerns the BsSMC complex ^24,50,52^. Previous studies concluded that BsSMC is recruited to the origin of replication through interactions with ParB protein ^50,52,57^, whereupon it progressively “zips” the left and right chromosome arms together along the entire length of the chromosome ^9,24,25^ (Fig. 3A). We constructed a strain containing an origin label (ParB-mScarlet), DNA label (HbsU-mTurqoise2), and BsSMC label (BsSMC-mGFP) in order to co-visualize the chromosome along with ParB and BsSMC (Fig 3B, Table S1).

**Figure 3.**
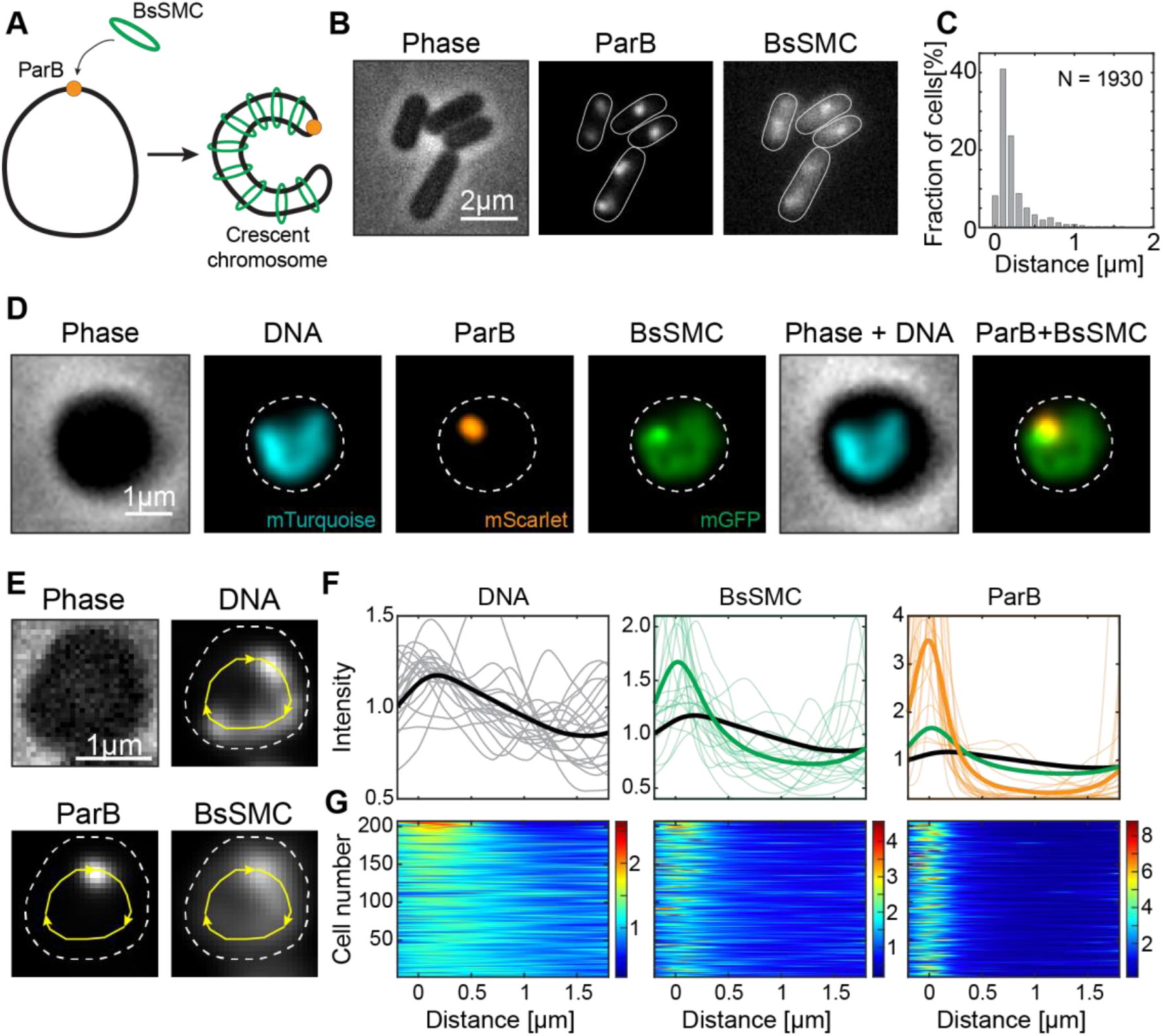
BsSMC proteins spread along the entire chromosome. **A)** Graphical representation of the crescent chromosome and the BsSMC positioning (cf. Wang *et al^1^*). **B)** Phase (left) and fluorescent (right) images of BSG4623 strain (*hbsu-mTurqoise2; smc-egfp; parB-mScarlet; P_hyperspank_-sirA*) after replication halt with 2 mM IPTG. **C)** Histogram of the distance between the BsSMC and ParB foci. The average distance between spots was d_avg_ = 126 ± 28 nm (mean ± std, n=1930). **D)** Images of bacterial strain BSG4623 strain after lysozyme (400 μg/ml) treatment for 30 min. **E)** Contour analysis of DNA, BsSMC, and ParB signals along the crescent chromosomes (cf. Fig. 2G.) **F)** Fluorescence intensity along the contour line of the chromosome, starting from the ParB locus: DNA signal (left, black line); BsSMC (middle, green); ParB (right, orange) (n = 215). **G)** Corresponding density plots along the chromosome in all individual cells. Colorbars represent the fold-increase.

We observed that SMC proteins positioned as a combination of typically 1-2 fluorescent foci with an additional clear signal that colocalized with the DNA signal over the entire chromosome (Fig. 3B). This starkly contrasts data for the ParB protein that showed that virtually all of the intracellular ParBs was captured into one focus, with very little background in the rest of the cell (Fig. 2B, D, S7)^58^. We quantified the distance of BsSMC-mGFP and ParB-mScarlet foci in rod-shaped cells, and observed a close proximity between the foci, i.e. an average mutual distance of only 126 ± 28 nm (mean ± std, Fig. 3C). Upon performing the same procedure in widened cells, we again observed the HbsU-labelled chromosome to adopt a crescent shape, with a ParB-mScarlet focus at the tip of the crescent-shaped chromosome (Fig. 3D). BsSMC-mGFP appeared to spread over the entire chromosome, with the highest intensity proximal to the ParB-mScarlet focus (Fig. 3D). In a control strain containing a tag only on BsSMC and not on ParB and HbsU, we observed the same distribution of BsSMC signal (Fig. S8).

We quantified the intensities of all signals along the contour of the chromosome (Fig. 3E) similar to those described in Fig. 2G-I. The average DNA signal, from HbsU-mTurquoise2, showed an increased condensation at the origin and a gradual decrease towards the terminus (Fig. 3F, G – left). The BsSMC-mGFP intensity, however, did not linearly scale with the DNA signal, but rather had a 1.6-fold increase at the origin compared to the average signal along the chromosome (Fig. 3F, G - middle), while the signal also gradually decreased towards the *ter*. A decreasing signal from origin to terminus can be expected if BsSMC is loaded at *ori* by ParB^50,52,57^ and removed at the *ter* by XerD^59^, while BsSMC exhibits a finite rate to dissociate from the DNA. Such a distribution was captured previously by ChIP-seq^60^, which was interpreted as stochastic unloading of BsSMC proteins along the chromosome arms^9,60^. In some cases, a secondary BsSMC focus was visible that did not correlate with the ParB location nor with an increased DNA density (Fig. 2G, S8). Finally, the ParB signal showed an even more pronounced peak near the tip of the chromosome and showed close to zero intensity signal along the rest of the contour length (Fig. 3F, G – right).

### Disruption of the BsSMC complex opens the crescent-shaped chromosome into a toroidal shape

Since BsSMC has been postulated to serve as the main connection between the left and right chromosome arms ^9,24,25^, we next tested for a potential reshaping of the chromosome after disruption of the BsSMC complex. For this, we constructed a strain with HbsU-mGFP as the chromosome label but included previously described ScpA-TEV3^61^ as well as *Pxyl-TEVp* (Table S1). ScpA is the kleisin subunit of the BsSMC complex that is a basic part of its ring-like structure, and it is essential in cells for fast growth ^62^. The incorporated Tobacco Etch Virus (TEV) protease recognition site (TEV3) was within the ScpA protein (Fig. 4A), while the TEV protease itself was included at a different locus under a xylose-inducible promotor. This allowed us to controllably disrupt the BsSMC complex by opening the SMC-kleisin ring upon xylose addition, and subsequently observe the chromosome re-shaping (Fig. 4A). We demonstrated high specificity of SMC disruption that only occurred in the presence of both the TEV3 cleavage site and xylose (Fig. 4B).

**Figure 4.**
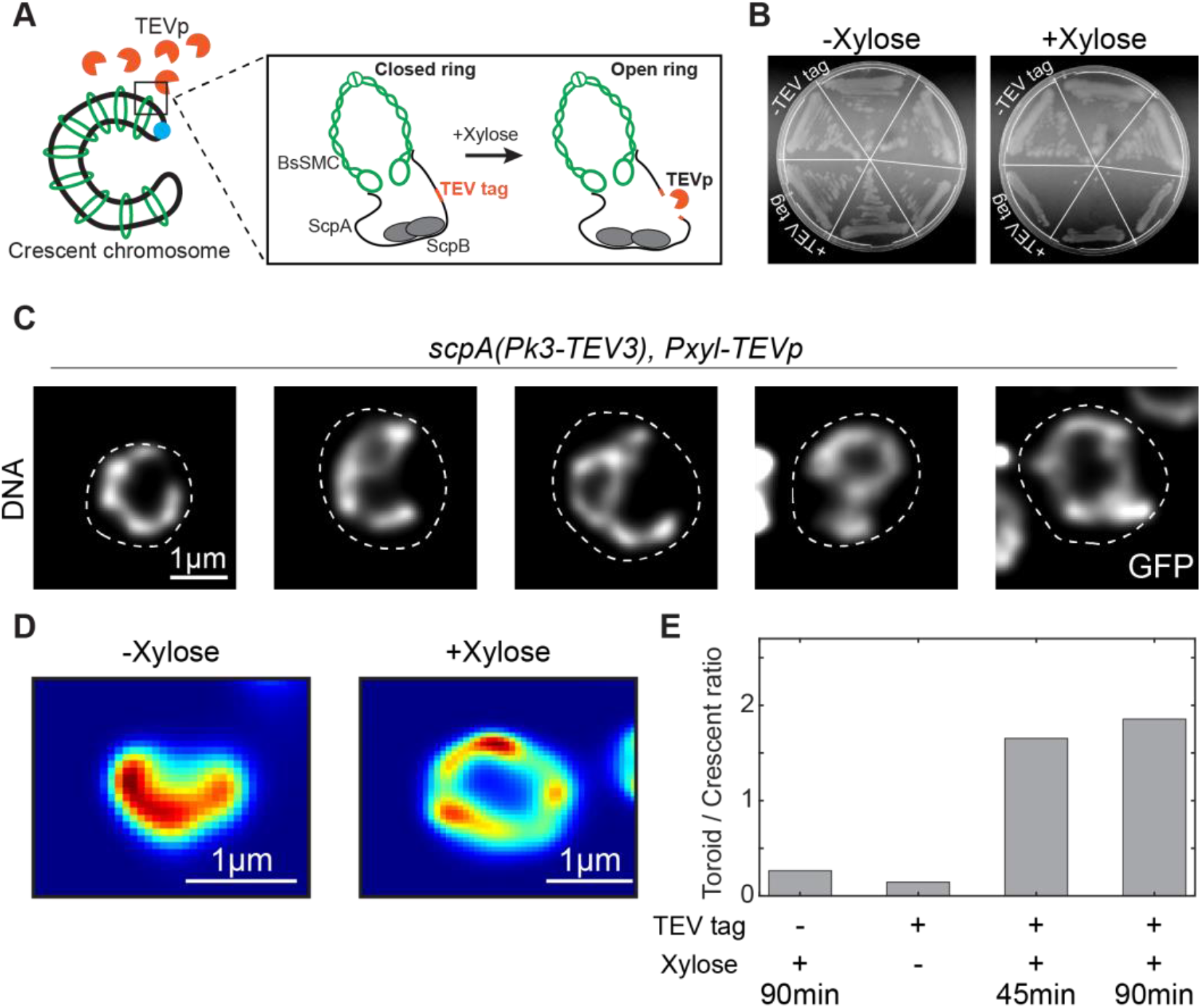
Disruption of BsSMC leads to the opening of the chromosome into a toroidal shape. **A)** Graphical representation of BsSMC along the crescent-shape chromosome in strain BSG219 (*dnaB(ts-134); hbsu-gfp; scpA(Pk3-tev3); P_xyl_-TEVp*). Zoomed region details the effect of the TEV protease that cuts the TEV cleavage site within the ScpA protein, which opens the BsSMC ring structure that supposedly is holding together the chromosome arms. **B)** Plating assay of strain BSG219 (depicted as +TEV) and its control BSG217 (depicted as -TEV) that is lacking the TEV cleavage site, in the absence (left) and presence (right) of 0.5% Xylose in the agar medium. **C)** Selected examples of the disrupted chromosomes in strain BSG219 after 60 min of xylose (0.5%) in the liquid medium. **D)** Representative examples of the crescent and toroidal chromosome shape with (right) and without (left) 0.5% xylose treatment. **E)** Relative ratios of toroidal to crescent chromosomes in four different conditions. See SI Fig. S10 and Methods for details on these estimates.

Expression of the protease resulted in dramatic macro-scale changes in the chromosome shape (Fig. 4C). Upon disruption of the BsSMC ring, the arms of crescent-shaped *B. subtilis* chromosomes opened. Sometimes, both arms separated entirely, resulting in a toroidal-shaped chromosome (Fig. 4C, S9). We observed that megabase-sized structural rearrangements of the chromosome occurred on the timescale of minutes, going from a well-defined crescent to a fully open state within ~30 min (Fig. S9A-B, Movie S3-S4). These changes were never observed in the absence of xylose, nor in a control strain lacking the ScpA-TEV3 recognition site (Fig. 4D-E, and Fig. S9C-D, S10). These results show that BsSMC proteins are indeed the sole agents that link the two arms of the circular chromosome of *B. subtilis*. The data are also in line with the previously reported chromosome rearrangements upon BsSMC deletion or ParB deletion^9,24,25^, whereupon chromosomes appeared to lose the second diagonal over time in ensemble Hi-C maps.

## Discussion

Direct observation of bacterial chromosomes can reveal novel insights into the underlying organization and dynamics of the DNA within the nucleoid. In *E. coli*, for example, direct microscopy observation of chromosomes in expanded-volume cells did resolve the local DNA condensation in the left and right chromosomal arm as well as a lack of structure within the *ter* domain ^16^. The approach furthermore yielded new insight into chromosome replication ^27,30^ and the role of SMC/MukBEF proteins ^15,28^ in chromosome organization.

Here, we applied a modified combination of cell-volume expansion with chromosome-copy control in *B. subtilis* to directly resolve the structure of the chromosome with single-cell microscopy. This showed that the chromosome exhibits a crescent shape with a non-uniform DNA density that is increased near the origin of replication. Our work expands on previous efforts by the Nollmann and Koszul labs who used Hi-C in combination with super-resolution microscopy to resolve the chromosome structure including its underlying domains^9^. They similarly observed that the replication origin contained a dense DNA region that they termed High-Density Region (HDR). Hi-C analysis showed a ‘second diagonal’, representing a juxtaposition of left and right chromosomal arms, and 3D modelling based on the Hi-C maps suggested a highly condensed origin of the replication. Due to the high compaction of the chromosome and high local density of the DNA, their 3D-SIM microscopy could not fully resolve the actual structure in the WT rod-shaped cells. Notably, their modeling suggested a helicoidal S-shaped or C-shaped 3D chromosome organization ^9^. Our live-cell observations of crescent-shaped chromosomes are very well compatible with these proposed models (Fig. 1, Movie. S1, S3).

Our data also revealed an increased DNA density in the origin region at the tip of the crescent-shaped chromosome, which may arise from multiple underlying biological processes. First, ParB proteins were proposed to condense the DNA near the *ori* site and form a partition complex by bringing multiple *parS* sites during the initial steps of chromosome segregation^9,48,63,64^. Second, ParB proteins have been proposed to recruit BsSMC proteins to the origin of the chromosome where a juxtaposition between the chromosomal arms is initiated ^24,50,52,57^. As our data do show an increased BsSMC content near the origin of replication, higher DNA density regions may result from an increased frequency of chromosome folding by BsSMC proteins. Third, the origin-of-replication genomic section contains highly transcribed genes in most bacteria, including *B. subtilis* ^65^. These genes are often accompanied by a high degree of supercoiling and plectoneme formation. This can further increase DNA condensation within the origin region compared to the rest of the chromosome. Maintaining only the origin region in a condensed state could have important implications for chromosome segregations via entropic forces ^66,67^; because highly condensed regions will be preferentially pushed towards the cell periphery in a cylindrical cell ^67^. Accordingly, deletions of BsSMC and ParB proteins mostly result in delayed origin segregation ^61,68^ and aberrant chromosome segregation in *B. subtilis* ^50,52^.

Our data also revealed an enrichment of BsSMC at the origin region (Fig. 3). This is in agreement with ChIP-seq data ^9,24,25,69^ and indicates that only a minor fraction of the total BsSMC on the chromosome are actively zipping the chromosomal arms, while a large fraction resides at the origin close to ParB. Although a recent biochemical study points to the direct interaction between ParB and BsSMC proteins^57^, the precise mechanism of BsSMC recruitment to the origin is not yet fully resolved.

Acute knock-down of BsSMC complexes resulted in large-scale chromosome reorganization where the left and right chromosomal arms lost proximity and the chromosome opened into a toroidal shape (Fig. 4, S9)^9,25^ that is reminiscent of the chromosome shape in *E. coli* cells^16^. In *E. coli*, the left and right chromosome arms showed a higher DNA condensation than the *ori* and *ter* regions, and it is this higher DNA density of the arms that was proposed to drive chromosome segregation via entropic forces ^67^. In *B. subtilis*, a knockdown of BsSMC proteins yielded a toroidal-shaped chromosome but it also resulted in aberrative segregation of nascent ParB partition complexes ^68^ and consequently aberrative chromosome segregation. Instead, DNA segregation in *B. subtilis* is mainly driven by the ParAB*S* system ^70^. These two model bacterial species thus employ very different strategies for enabling proper chromosomal segregation within daughter cells.

Our study investigated the *B. subtilis* chromosome organization via direct live-cell imaging. The data revealed the chromosome shape and distribution of ParB and BsSMC proteins within single cells, and we showed a disruption of chromosome shape in single cells upon BsSMC knock-down. We observed that the origin of the replication is maintained in a condensed state even under non-confining conditions. The cell-shape manipulation imaging approach can be applied to other bacteria, allowing for single-cell real-time imaging of the chromosome and the dynamics of intercellular processes.

