## Supplementary information and figures for "Direct observation of a crescent-shape chromosome in *Bacillus subtilis*"

**Abstract**

Bacterial chromosomes are folded into tightly regulated three-dimensional structures to ensure proper transcription, replication, and segregation of the genomic information. Direct visualization of the chromosomal shape within bacterial cells is hampered by cell-wall confinement and the optical diffraction limit. Here, we combine cell-shape manipulation strategies, high-resolution fluorescence microscopy techniques, and genetic engineering to visualize the intrinsic shape of the bacterial chromosome in real-time in live *Bacillus subtilis* cells. We show that the chromosome exhibits a crescent shape with a non-uniform DNA density that is increased near the origin of replication (*oriC*). Additionally, we localized ParB and BsSMC proteins – the key drivers of chromosomal organization – along the contour of the crescent chromosome, showing the highest density near *oriC*. Opening of the BsSMC ring complex disrupted the crescent chromosome shape and instead yielded a torus shape. These findings help to understand the threedimensional organization of the chromosome and the main protein complexes that underlie its structure.

### Methods

#### Strain construction

All strains were constructed using the homologous recombination-based cloning described in detail in Diebond-Durant et al[71]. In brief, we used a transformation of recombinant DNA to create *B. subtilis* strains at the *smc, scpA, parB, amyE, hbsu* and *thrC* loci by allelic replacement harboring natural competence induced by the stationary phase growth as well as starvation due to media depletion (1-2 h incubation in SMM medium lacking amino acid supplements) the same as described in detail Diebold-Durand et al[71]. We selected the new strains on SMG-agar plates in the presence of the appropriate antibiotic at 30°C for 16 h. We verified the correct genotype of single colonies by antibiotic resistance profiling on solid agar plates, colony PCR, and locus sequencing and stored the correct clones at -80°C.

#### Bacterial growth conditions

Prior to all experiments, bacterial strains stored at -80°C, were streaked on a 1xNA (Oxoid) agar plate (1.5% agarose). Plates were incubated at 30°C for 10-16 h, overnight to obtain single colonies, and used for a maximum of 2 days after that. One day before the experiments we picked single colonies and incubated them in 5 ml liquid osmoprotective medium for 10-16 h, overnight. Osmoprotective media (from now also referred to as SMM+MSM) used in this study were modified from Kawai et al. [36] and composed of 2×MSM (40 mM magnesium chloride, 0.2 M-1 M succinate, 40 mM maleic acid, 0.2% yeast extract) mixed in 1:1 relation with 2xSMM (0.04% magnesium sulphate, 0.2% sodium citrate dehydrate, 0.4% ammonium sulphate, 1.2% potassium dihydrogen phosphate, 2.8% dipotassium phosphate) and supplemented with 0.2% L-tryptophan, 1 μg/ml ferric ammonium citrate, 0.1% glutamic acid, 0.02% casamino acids (Bacto™), 1 mg/ml L-threonine. We supplemented the overnight cultures with 5 µg/ml kanamycin, 1 µg/ml erythromycin, 5 µg/ml chloramphenicol, 100 µg/ml spectinomycin where applicable (see Table S1), but the cultures on the day of experiments were not exposed to selection antibiotics. We made 2xMSM solutions with the following range of succinate concentrations 0.2 M, 0.4 M, 0.6 M, 0.8 M, 1 M, for the purpose of testing osmoprotective conditions with the sugar concentration ranging from 100-500 mM (see Fig. S1). On the day of the imaging experiments, we diluted the overnight cultures 50x in the total of 10 ml osmoprotective media. We grew the cultures for 3 h (30°C, 200rpm orbital shaking), until they reached the early exponential phase, and then induced replication halt either by supplementing with 2 mM IPTG or moving them to 37°C and 200rpm orbital shaking, depending on the strain, for a total of 30-180 min depending on the experiment. In the case of TEV protease expression experiments using the strains BSG217 and BSG219 (Table S1), we added 1.5% xylose 45 min before imaging for timecourse imaging (Fig. 4). For the real-time imaging of BsSMC degradation we included the xylose (at 1.5% final concentrations) into the agar pad just before imaging.

**Conversion to L-form spheroid cells**

After growing the bacteria for a targeted time of replication halting, we subjected the shaking bacterial cultures to lysozyme treatment. We made a 100 mg/ml fresh stock of lysozyme on the day of the experiment by mixing Lysozyme from chicken egg white (Sigma-Aldrich) with 1xPBS (140 mM NaCl, 10 mM phosphate buffer, and 3 mM KCl, pH 7.4, Phosphate Saline Buffer, (Sigma-Aldrich)) and kept it on ice for the duration of that day (4-6 h). Depending on the temperature of the cell growth we added the lysozyme to a final concentration of 400 µg/ml either 20 min, when grown at 37°C, or 30 min before, when grown at 30°C.

**Fluorescence imaging**

After cell growth and necessary treatments in liquid culture, we transferred ~3 ml of the liquid cell culture to the 35 mm glass bottom dish (MatTek), which we beforehand passivated with 100 µg/ml UltraPure$^{TM}$ BSA (Invitrogen) for 15-20 min. We specifically avoided using any pipetting steps after the lysozyme treatment to maintain the integrity of spheroidal cells and prevent them from lysis due to shear forces. We then centrifuged the imaging dishes custom-made holders at 400g for 4 min in a swingout centrifuge with 96-well plate holders (Centrifuge 5430 R, Eppendorf). We removed residual supernatant containing non-adherent cells by pipetting at the edge of the dish and placed an agarose pad (~ 4 x 4 mm) containing 1x osmoprotective medium (lacking yeast extract) and 0.2% low melting agarose (Promega, Madison, USA) on top of the glass bottom. Note that yeast extract needed to be removed from the medium used for making the soft imaging agar pad due to residual background fluorescence in the green channel (488 nm). We placed a small 11x11 mm glass coverslip on top of the agar pad to assure the cells were not floating in the medium during imaging due to liquidity of 0.2% agarose and to prevent premature evaporation and drift.

We carried out widefield Z-scans out using a Nikon Ti-E microscope with a 100X CFI Plan Apo Lambda Oil objective (NA=1.45). The imaging stage was maintained at 37°C for experiments using temperature-sensitive mutants, and at room temperature for mutants affected by IPTG expressing of SirA protein. We excited DAPI using SpectraX LED (Lumencor) filter cube with $\lambda_{ex}$/ $\lambda_{bs}$/ $\lambda_{em}$ = 363–391/425/435–438 nm, GFP and SYTOXGreen using filter cube with $\lambda_{ex}$/ $\lambda_{bs}$/ $\lambda_{em}$ = 485-491 nm/506 nm/501-1100 nm, and mScarlet and STYOXOrange using filter cube with $\lambda_{ex}$/ $\lambda_{bs}$/ $\lambda_{em}$ = 540–580/585/592–668 nm. We used an Andor Zyla USB3.0 CMOS Camera to collect the fluorescent signals. We took 9-11 Z-slices of step size l = 200 nm, which accounts for a total of 1.8-2.2 µm of scanning volume, that we fed into the deconvolution software (see below).

**Solid agar plating assays**

For testing the growth differences in presence/absence of xylose in the strains carrying *scpA(Pk3-TEV3)::specR* locus (Fig. 4B), we used a standard LB-agar (1.5% agarose). As described above, we grew the strains on separate plates carrying the selective antibiotic (here 5 µg/ml chloramphenicol and 100 µg/ml spectinomycin) one day before the experiment. We picked three single colonies from each plate and re-streaked them in a radial orientation on the plate containing LB-agar or LB-agar + 1.5% xylose. The plates were left to incubate at 30°C for 16 h, after which we imaged them on a Gel imager (BIO-RAD Chemidoc™).

**Growth curves**

For monitoring bulk growth (Fig. S6), we first grew the strains in the selective media overnight, as mentioned previously. On the day of the experiment, we diluted the overnight cultures 40x in the fresh SMM+MSM medium and grew them until the OD reached ~0.2 - 0.3. We then again diluted the cultures to the final OD of 0.01 and distributed in 96-well plate (Nunc), with the final volume of 170 µl. The plates were loaded into an Infinite 200Pro fluorescence plate reader (Tecan, Männedorf, Switzerland) and incubated at 30°C with the orbital shaking (2.5 mm amplitude) for a period of 24 - 48 h. Cell density was measured at 600 nm at 15 min intervals. We performed all experiments using at least biological triplicates, and in some instances more.

**Deconvolution microscopy**

For deconvolution microscopy we used Huygens Professional deconvolution software (Scientific Volume Imaging, Hilversum, The Netherlands), using an iterative Classic Maximum Likelihood Estimate (CMLE) algorithm. We measured the point spread function

(PSF) experimentally by using 200 nm Tetraspeck beads (Invitrogen) and the recommended Huygens Professional guidelines (*https://svi.nl/Point-Spread-Function-(PSF)*). Stacks of 9-11 Z-slices were fed into the Huygens Professional software and we deconvolved each signal channel separately.

**Structured illumination microscopy imaging**

For SIM microscopy, samples were prepared as previously described, except using two rounds of centrifugation with 3 ml cell culture in order to obtain higher cell density to accommodate for the smaller camera FOV. We used a Nikon Ti-E microscope with an AiRSIM module and a 100X CFI Apo Oil objective (NA=1.49). We used a 3D-SIM option with the 5 pos x 3 angles for imaging our live samples, and a Z-stack of 11 positions with l = 200 nm in between. NIS-Elements (version 5.2.1) software was used for image reconstruction where we carefully adhered to the Nikon N-SIM reconstruction guidelines and parameters to avoid reconstruction artefacts (Fig. S5).

**Image processing and analysis**

For obtaining the cell outlines and counting fluorescent foci within each cell, as well as for measuring distances between ParB-mScarlet and BsSMC-GFP foci within each cell, we used the image segmentation and analysis software Oufti[72]. We obtained subpixel precision *B. subtilis* cell outlines using the cellDetection tool in Oufti with the parameters shown in Table S2. We used the detectSignal tool for fluorescent foci detection of the ParB-mScarlet and BsSMC-GFP signals using the selection parameters shown in Table S2.

We used custom-written MATLAB scripts, modified from the original publication of Kaljevic *et al.*[73], to extract the fluorescent foci positions, count the number of foci per cell, and determine the distances between foci in each channel. These scripts are open-source and available online (doi.org/10.5281/zenodo.7615509).

For a systematic and detailed analysis of the chromosome shape, shown in e.g. Fig. 2 and Fig. 3, we used home-developed Matlab code based on earlier work from Wu et al[16]. The image processing covered the following elements:

*Selection* - First, we used Fiji/ImageJ to user-select ROIs around round cells based on the phase image alone, yielding typically ~$10^2$ cells per field-of-view (FOV). This ROI list was imported in Matlab. For each ROI, we obtained a mask outlining the cell wall from the wide field image. Within this cell mask, we analysed the signals of the other colour labels, depicting the patterns representing DNA, SMC, and ParB.

*Screening* - For a randomly chosen FOV, the selected round cells displayed a large variety of patterns and artefacts and sometimes included misdetections due to the thresholding. The latter were screened out by a simple cell roundness criterion. Furthermore, multiple ParB spots could lie in different focal planes such that only one was observed per focal plane. To make sure we selected single chromosomes, we applied a projection of maximum intensity along the defocus stack and accepted only those cells that had a single ParB spot appearing in maximum projection.

*Backbone pattern analysis* - Here, we made use of the characteristic shape of the chromosome pattern. As seen in-plane, the patterns formed crescent arcs that broadly followed the inside cell wall over a broad annular section of 90-180 degrees. As described before, such an arc-like structure could well be described in an annular coordinate system, where a 'backbone' centre line runs over the chromosome arc[16]. Using this backbone as a length axis, we sampled the relative intensity of the respective labels per unit annular section along this length. To compare many cells, we performed this sampling over a fixed, representative length, starting at the angle where the ParB signal maximizes, since this maximum represents the location of the *ori* site on the chromosome. We thus created intensity vs distance profiles for all labels. Notably, the cells with the crescent-shape chromosome may appear in a regular fashion or lie upside down. Therefore, starting from the *ori* location, the crescent may point clockwise or counter clockwise. To compare all cells, we flipped the profiles such that for all, the crescent intensity moved towards positive length values. We collected all these profiles in demographs shown in Fig. 2I and Fig. 3G. Each line in a demograph represents a heatmap of the DNA density profile from the origin (position of 0μm) to the end of the chromosome. These lines are stacked for all cells (y-axis) to make the common trends well visible.

*Foci and cluster analysis* - Following an earlier described method[16], we described the measured intensity patterns as a sum of clusters. Here, the chromosome pattern was described as a group of separate clusters. Each of these clusters was then described with a sub-group of psf-limited spots. For this sub-group, any of the spots was within one psf of at least one other. This resuted in a smooth, compact shape of this cluster. On the other hand, all spots in another cluster would be further away than one psf, so that there was always a distinguishable gap between each two neighboring clusters. In this way, the sometimes complicated 'blob patterns' can be described in a compact way as a limited set (typically, 4-5 per cell) of optically separable clusters, with each of these clusters coming with a particular xy position and a relative intensity (as compared with the intensity counted in the entire cell area). In the case of a compact focus, the cluster description simplifies in just one psf-sized spot. If a single cluster extended beyond one psf, such as the somewhat smeared-out spots observed for the ParB label, this approach still provided us with an intuitive description of the amount of fluorescence associated with one ParB spot. Finally, the xy positions of the cluster analysis can be correlated to the above-described backbone length axis, allowing to readily and quantitatively compare the relative intensities of mutually associated clusters of DNA, ParB and SMC (Fig. 3).

**Chromosome shape selection**

We were unsuccessful in obtaining the ParB-mScarlet tag in strains BSG217 and BSG219, containing HbsU-tag, $P_{xyl}$-*TEV*p and/or ScpA-TEV3 in multiple cloning attempts. While we cannot distinguish between single and partially replicated chromosomes, due to strain construction issues, we attempted to quantitatively distinguish between torus-shaped chromosomes and crescent-shaped chromosomes in the presence of 0.5% xylose. For this, we performed the following manual user selection. We collected one image from four different samples (Fig. 4E, S10), which describe four experiments with different expected outcomes. Each of these was intensity thresholded to detect ROIs with separated DNA patterns based only on the continuous fluorescent signal. Next, a randomly picked ROI, containing a signal from a single cell, was presented to a user for classification of this pattern. The user could choose from one of four categories (toroid, crescent, compact, other/undefined) without knowing from which original image the ROI was taken. This was done for a series of 100 picks each, with no ROIs being presented to the user more than once within this series. To obtain an estimate for sampling variation, we repeated this series 20 times per user for a total of 2000 selected ROIs. To evaluate possible user bias, the procedure was done independently by two users (MT and JK) shown in Fig. S10. We then traced back the described ROIs (in one of those four categories) to the original images and plotted the selection distribution shown in Fig. 4E and Fig. S10.

**Visualization and representation**

All fluorescent and phase images were visualized in the open-source image-analysis software Fiji[74]. We used a calculated microscopic pixel size of 65.35 nm/px in all images for scale bar creation. Adobe Illustrator 2020 was used for visualization and manuscript figures. After reconstructing the SIM microscopy images, we used SIMcheck software package for visualization [75] (Fig. S5) and a Fiji in-built function "3D project" to create a 3D reconstruction of the crescent chromosome (Movie S1), as well as the plug-in 3D viewer to create the isosurface plots of the same image (Movie S2). The same was used to visualize the deconvolved 3D stacks in Movie S3-S4.

**Acknowledgments:** We thank Jaco van der Torre and Hammam Antar for useful discussions and advice during the project.

**Funding:** We acknowledge funding support for the work in CD lab by the European Research Council Advanced Grant 883684 as well as by project OCENW.GROOT.2019.012 which is financed by the Dutch Research Council (NWO). We also acknowledge funding for the work in SG lab by the Swiss National Science Foundation (grant number: 310030L_170242).

**Author contributions:** Conceptualization: MT, SG, CD. Cloning and strain construction: MT, FPB. Experimental work: MT. Formal analysis: MT, AJ, JK. Methodology: MT, FPB, JK. Visualization: MT. Funding acquisition: SG, CD. Supervision: SG, CD. Writing – original draft: MT, CD. Writing – review & editing: MT, FPB, JK, AJ, SG, CD.

**Competing interests:** Authors declare that they have no competing interests.

**Data and materials availability:** All data included in the manuscript are available in the main text or the supplementary material. All raw microscopy data is available upon request. The analysis scripts, including the test dataset for new users, are freely available in Zenodo open repository (doi.org/10.5281/zenodo.7615509).

| Name | Genotype | Origin |
|---|---|---|
| BSG001 | 168 ED, trpC2 | Gruber lab |
| BSG217 | 168 ED, dnaB(ts-134; K85E), ΔamyE::Hbsu-GFP::CAT, scpA::specR, thrC::Pxyl-TEVp::ermC, trpC2 | This study |
| BSG219 | 168 ED, dnaB(ts-134; K85E), ΔamyE::Hbsu-GFP::CAT, scpA(Pk3-TEV3)::specR, thrC::Pxyl-TEVp::ermC, trpC2 | This study |
| BSG1001 | 1A700, trpC2 | Gruber lab |
| BSG4595 | 1A700, ParB-mScarlet::kan, amyE::Phyperspank-opt.rbs-sirA (spec), trpC2 | This study |
| BSG4596 | 1A700, amyE::Phyperspank-opt.rbs-sirA (spec), trpC2 | This study |
| BSG4612 | 1A700, smc-mGFPmut1 ftsY::ermB, amyE::Phyperspank-opt.rbs-sirA (spec), trpC2 | This study |
| BSG4623 | 1A700, smc::-mGFP1mut1 ftsY::ermB, hbsU-mTorquais::CAT, ParB-mScarlet::kan, amyE::Phyperspank-opt.rbs-sirA (spec), trpC2 | This study |

**Table 1. Bacterial strains used in this study**

| Parameter name | Value |
|---|---|
| **cellDetection** | |
| ThreshFactorM | 0.9925 |
| ThreshMinLevel | 0.8 |
| EdgeSigmaL | 1 |
| **spotDetection** | |
| minHeight | 0.002 |
| minWidth | 0.5 |
| maxWidth | 10 |
| Adjusted Squared Error | 0.4 |
| **objectDetection** | |
| Bkg substraction method | 3 |
| Bkg substraction threshold | 0.1 |
| Bkg filter size | 8 |
| Smoothing range (pixels) | 3 |
| LOG filter | 0.1 |
| Sigma of PSF | 1.5 |
| Fraction of object in cell | 0.4 |
| Minimum object area | 50 |

**Table 2. Oufti detection parameters used in this study**

**Supplementary Figures S1-S10**

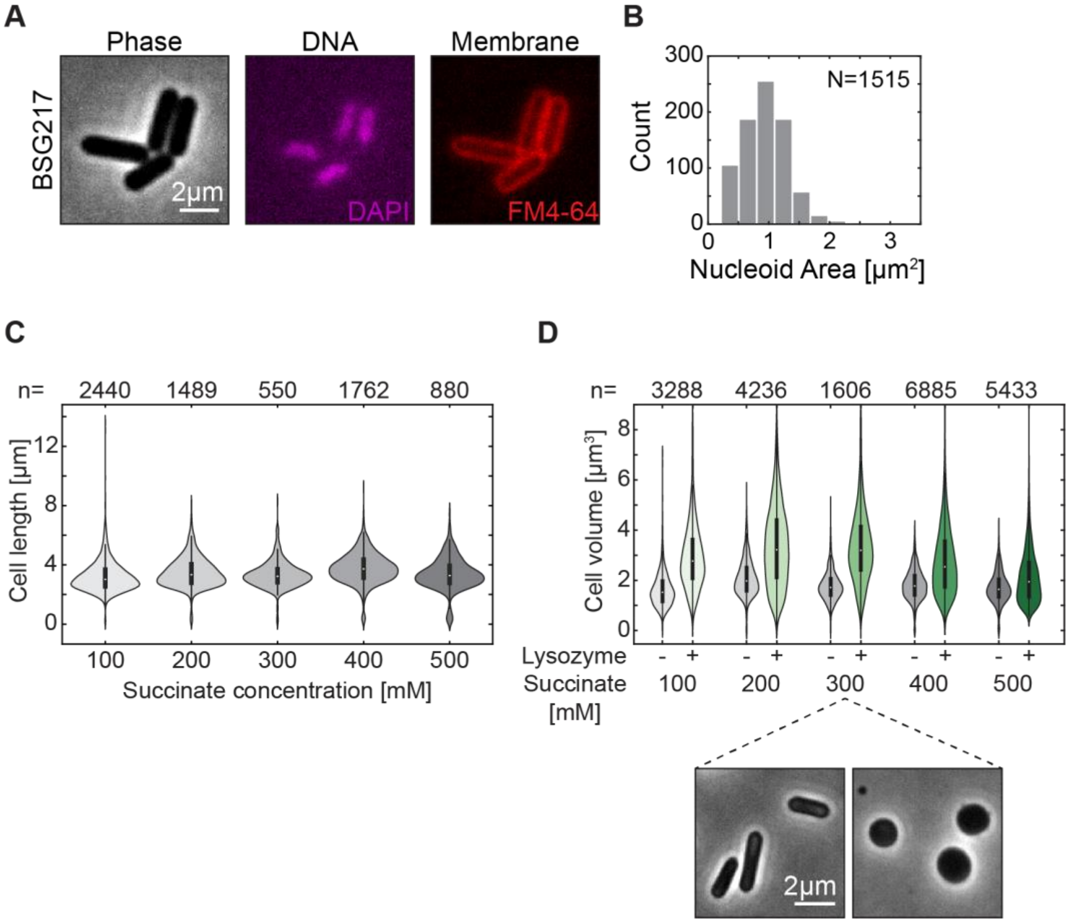

**Figure S1. Transformation from cylindrical to spherical cell shape leads to an increase in volume. A)** Phase and fluorescent images of the BSG217 strain (see Table S1), with labelled DNA and cell membrane, after replication halt at 37$^{o}$C for 60 min. **B)** Nucleoid area in replication halted cells shown in A). Mean nucleoid length and width were l=1.28 ± 0.49 μm and w=0.72 ± 0.09 μm (mean ± std, N = 1515). **C)** Longitudinal cell length of rod-shaped *Bacillus subtilis* cells (strain BSG217) grown in SMM+MSM medium of different osmolarities. **D)** Cell volumes of rod-shaped and spherical cells (see Methods and ref 2) grown in the same SMM+MSM medium of different osmolarities. Gray data represent samples that were not exposed to lysozyme treatments; green data represent cells exposed to lysozyme treatment (see Methods).

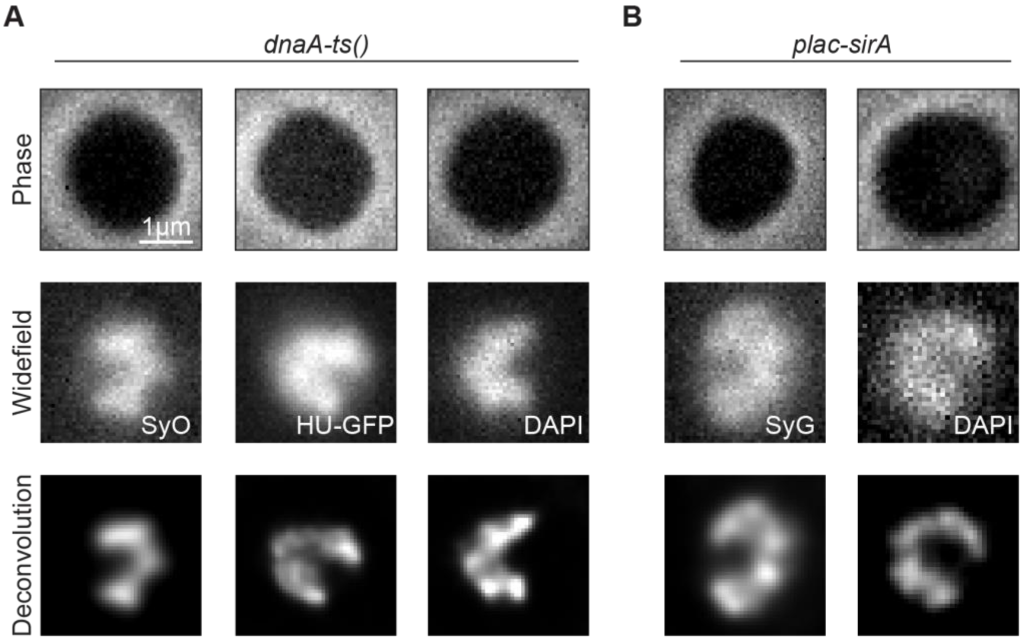

**Figure S2.** ***Bacillus subtilis*** **chromosome adopts a crescent shape regardless of replication halt strategy or DNA-visualization dye. A)** Phase and fluorescent images of crescent chromosomes in BSG217 cells after lysozyme treatment (400 µg/ml for 20 min) using different DNA dyes (SYTOXOrange (250 µg/ml), GFP-fusion, DAPI (3 µg/ml)). Top to bottom – phase image of spherical bacterial cells, widefield fluorescence image, and the same images after deconvolution via Huygens Professional software (see Methods). **B)** Same as A) for the BSG4595 strain that is replication halted using 2 mM IPTG for 90 min.

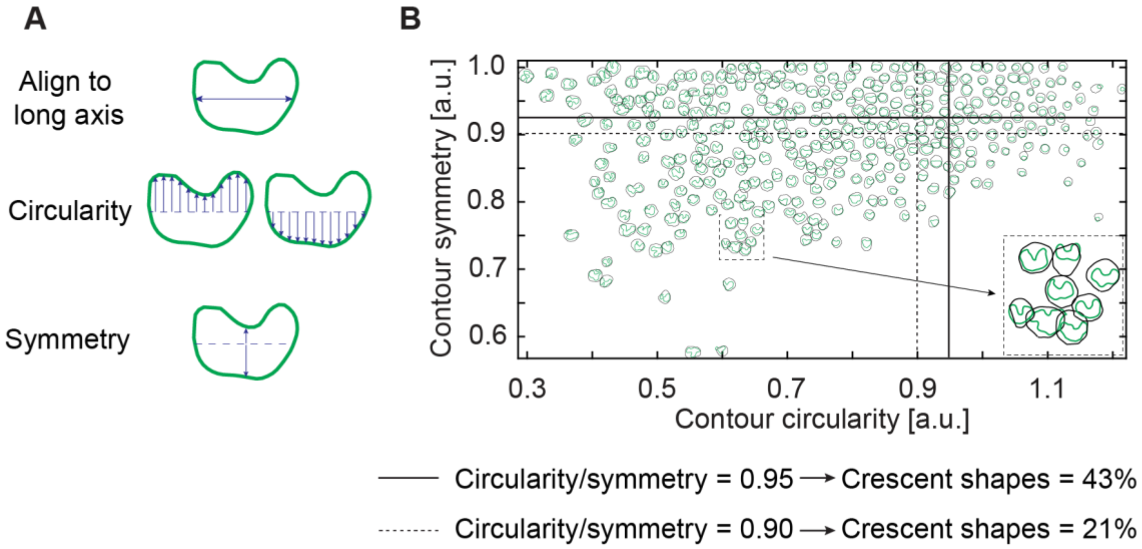

**Figure S3. Quantification of crescent-shaped chromosomes in spherical cells that were exposed to hypoosmotic medium. A)** Graphical representation of the stepwise selection of crescent chromosomes. Initially selected chromosomes as Oufti objects (see Methods) are positioned along their long axis. Then a ‘circularity estimator’ is applied that estimates the number of equidistant points in the top and bottom of the mid-axis. Finally, a ‘symmetry estimator’ measures the relative difference in the distance from the midpoint to the mid-axis. **B)** Cells plotted by their circularity and symmetry. Green line represents the Oufti object outline (which characterizes the chromosome contour)[2]. n = 1321. Selection at threshold of 0.90 (dashed line) or 0.95 (full line) for the ration of circularity/symmetry results in 21% or 43% crescent shapes, respectively.

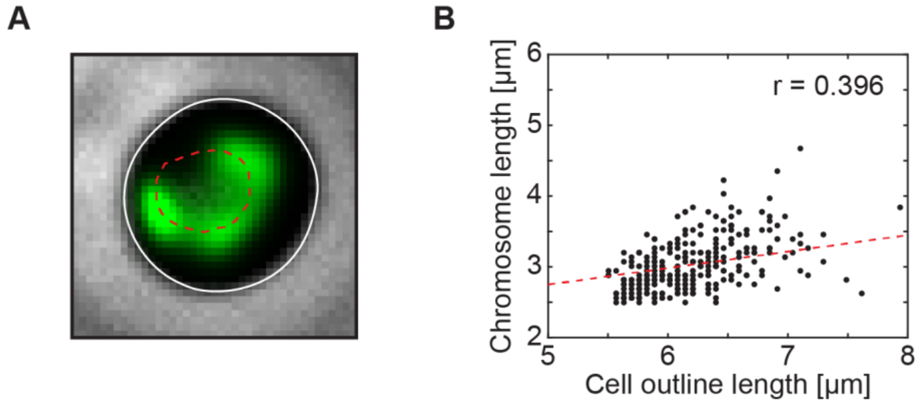

**Figure S4. Crescent chromosome full expansion is weakly correlated with final cell expansion. A)** The micrograph shows the cell outline in white full line, and crescent chromosome contour in red dashed line. **B)** Chromosome contour length versus cell-boundary contour length in spherical cells BSG4595. Black dots represent individual data points. N = 292, r = 0.3965.

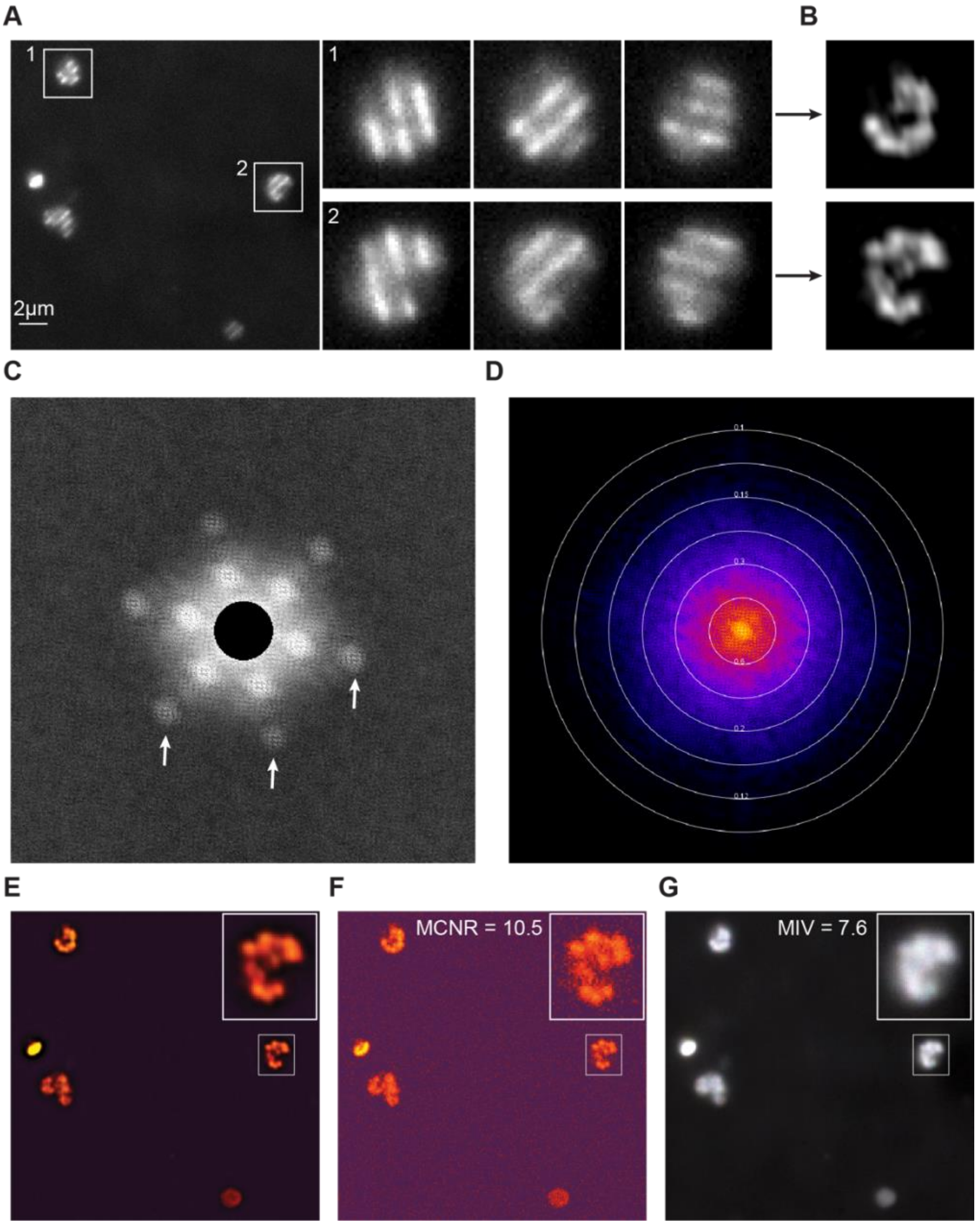

**Figure S5. Structured Illumination Microscopy image controls. A)** Individual frame from the raw image sequences of 5x3 imaging (position x angle) in the SIM microscope. Two cells of the strain BSG217 (see Table S1) are zoomed and shown with the raw image in the three frames. **B)** Reconstructed SIM image of these individual cells. **C)** Fourier projection of the raw data of the full image (in panel A) in reciprocal space. The image shows the first and second order point of high frequency in all the angles (white arrows), which we used as mandatory to pass the quality control. **D)** Fourier-space image of the reconstructed image overlaid with concentric circles showing corresponding the spatial resolution (in µm). Based on the image, we estimate a resolution of ~0.16 µm. **E)** Modulation Contrast map for reconstructed bottom image from panel A-B). **F)** Same as in E) but for raw image. The MCNR (modulation contrast-to-noise ratio) passes the SIMCheck quality control[3]. **G)** Motion and illumination variation (MIV) in different angles and frames. The gray image with an absence of colors indicates a high stability and low variation between angles during illumination in the SIM imaging, which ensures that the total imaging sequence for one image is faster than any visible movements of the *B. subtilis* chromosome.

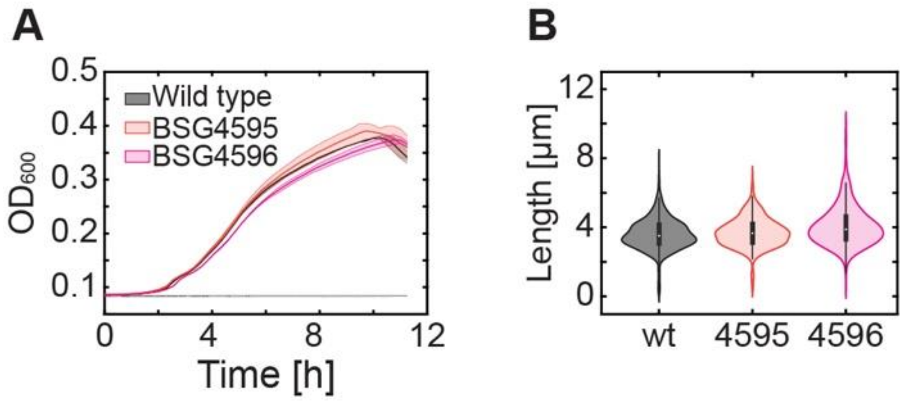

**Figure S6. Bacterial growth and phenotype is not affected by genetic edits to *parB* and *amyE* loci. A)** Tecan plate reader growth curves (see Methods) for B. subtilis 1A700 strain and BSG4596, BSG4595 strains, carrying $P_{lac}$-*sirA* and $P_{lac}$-*sirA* + *parB-mScarlet* (used in extensive quantification in Fig. 2), respectively. **B)** Longitudinal cell length in phase images for the wild-type strain and $P_{lac}$-*sirA* containing modified strains.

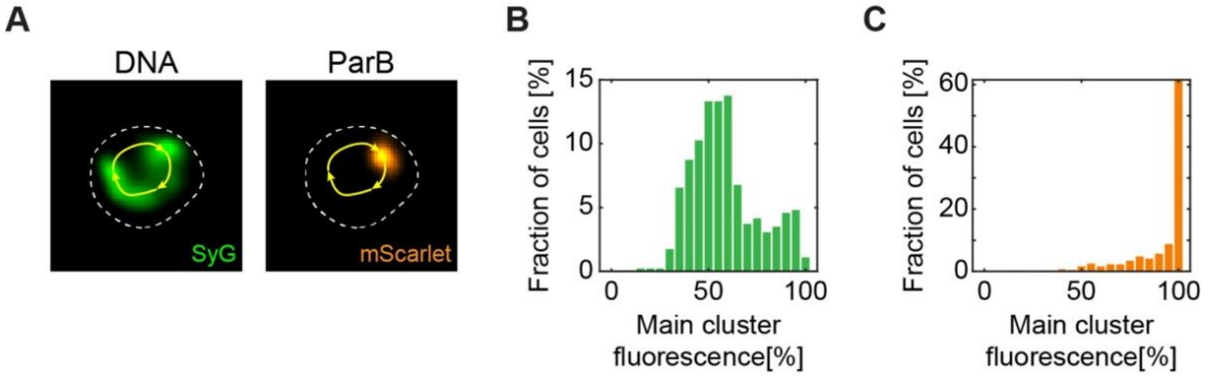

**Figure S7. DNA and ParB clusters adopt different fractions of the total signal. A)** Representative fluorescence images of the crescent chromosome and ParB focus (cf. Fig. 2) in the strain BSG4595. Dashed white line represents cell outline, and yellow arrows represent a sketch of the contour line measurement along the crescent chromosome. **B)** Relative DNA presence (based on fluorescent signal, see Methods) in the main cluster compared to the total DNA signal within the crescent chromosome. **C)** Same for the ParB signal.

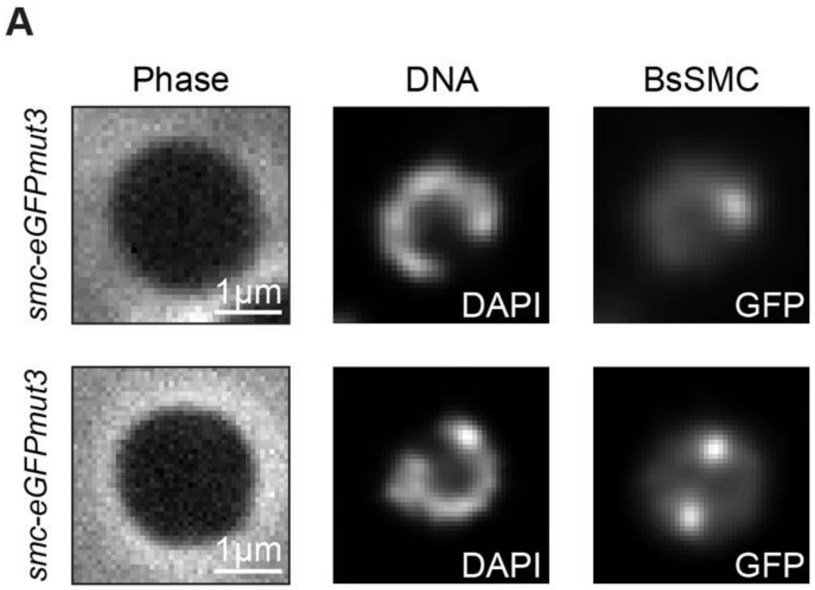

**Figure S8. BsSMC proteins localize along the contour of the crescent shape and form fluorescent foci along it. A)** Phase and fluorescence images of BSG4612 strain (see Table S1) containing $P_{lac}$-*sirA* and BsSMC-eGFPmut3 label. Top: example with a single BsSMC focus close to the tip of the crescent chromosome. Bottom: example with multiple BsSMC foci.

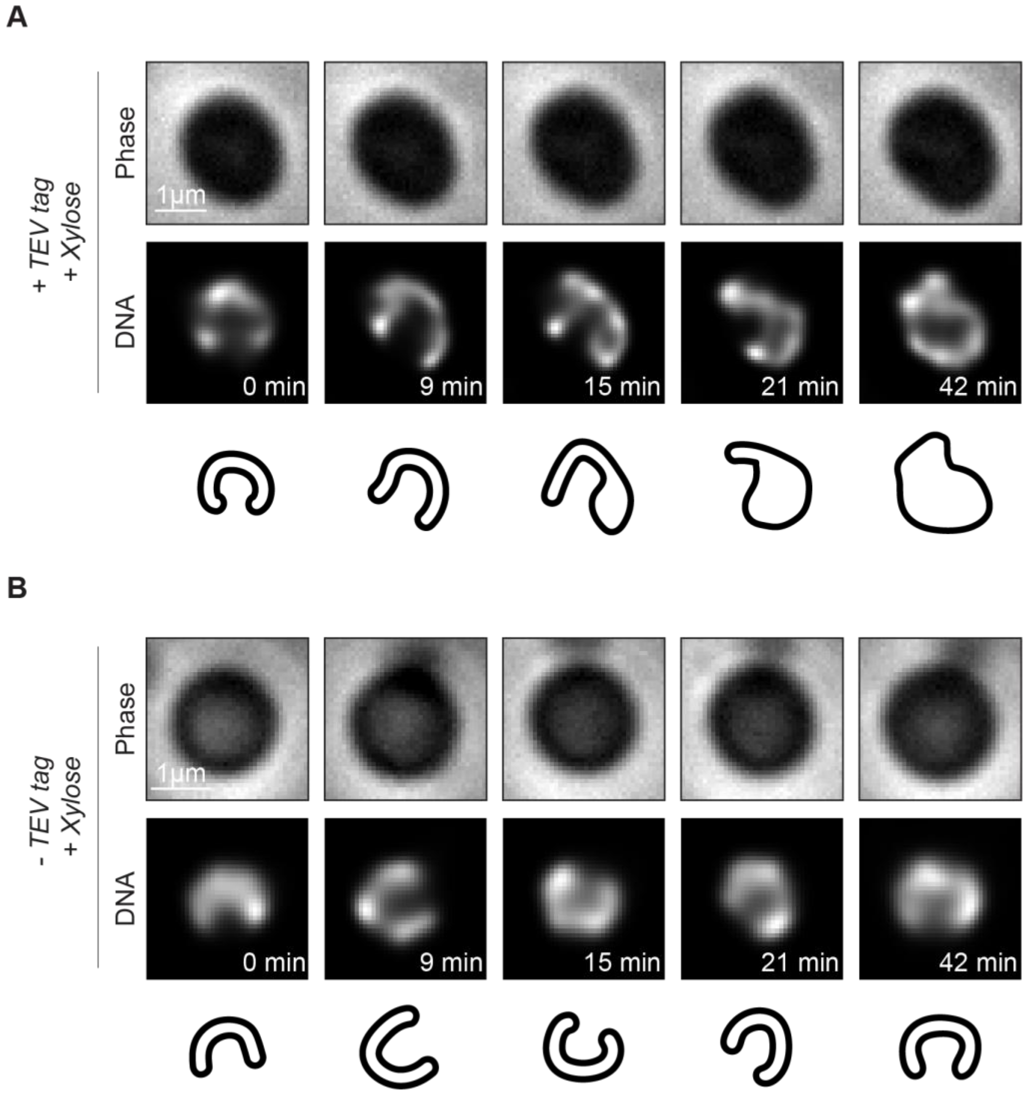

**Figure S9. Real-time imaging of BsSMC protein knock-down via xylose-expressible TEV protease shows a reshaping of the chromosome which loses its crescent shape. A)** Timelapse imaging ofa single *B. subtilis* **chromosome** in strain BSG219, containing the ScpA-TEV3 in presence of xylose-expressed wTEV protease (0.5% xylose). Schematic representation of chromosome shapes is represented below. **B)** Control experiment showing timelapse images under the same conditions as in A) but for strain BSG217 that does not contain the TEV tag and TEV protease.

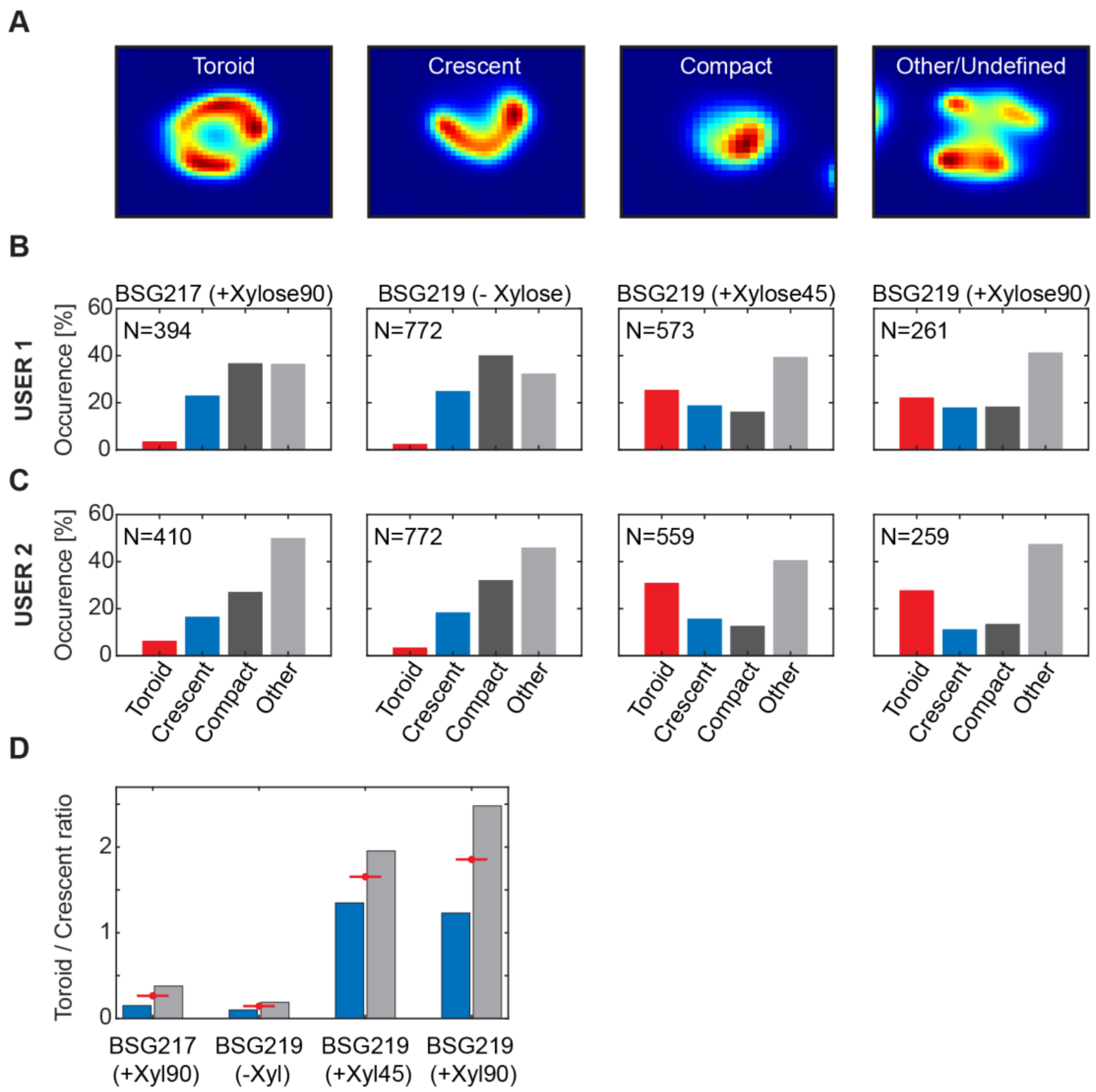

**Figure S10. Randomized identification of chromosome shapes under SMC knock-down conditions. A)** Example images representing four chromosome categories that were presented to two independent users for identification (see Methods for detailed description). **B)** Distribution of four selected categories over different samples (shown on top) for user 1 (MT). **C)** Same) for user 2 (JK). **D)** Ratio of torus-shaped to crescent-shaped chromosomes in all samples. Blue and grey bars represent the two users; mean values of the two users are represented by the red lines.
